# Dexmedetomidine produces more sleep-like brain activity compared to propofol

**DOI:** 10.1101/2025.05.27.656383

**Authors:** Bryan M. Krause, Emily R. Dappen, Rashmi N. Mueller, Hiroto Kawasaki, Robert D. Sanders, Kirill V. Nourski, Matthew I. Banks

## Abstract

**Introduction:** Dexmedetomidine is a selective α_2_-adrenergic agonist used as an anesthesia adjunct to produce a state of sleep-like sedation. However, how brain activity compares quantitatively during dexmedetomidine anesthesia to that during natural sleep, and thus just how “sleep-like” dexmedetomidine anesthesia is, remains unclear. Previously, we showed that the general anesthetic propofol is associated with changes in connectivity and cortical network structure comparable to those observed during sleep. Here, we compare the effects on brain activity of dexmedetomidine, propofol, and sleep quantitatively using intracranial encephalographic (iEEG) recordings in human research participants.

**Methods:** iEEG recordings were obtained in 34 epilepsy patients being evaluated for potential seizure resection surgery. Band power and functional connectivity (alpha weighted phase lag index, gamma envelope correlations) and network entropy were measured in recordings during task-free (“resting state”) periods just prior to surgery during anesthesia with either dexmedetomidine or propofol, and during overnight sleep. Anesthesia stage (wake, sedated, unresponsive) was determined using the Observer’s Assessment of Arousal and Sedation. Sleep was staged using standard polysomnography.

**Results:** As expected, significant differences in delta power were observed during dexmedetomidine and propofol as well as during sleep. However, the magnitude of changes in delta power were smaller and regionally heterogeneous for propofol compared to dexmedetomidine and sleep. Functional connectivity changes were comparable between dexmedetomidine, propofol, and natural sleep. Significant changes in network entropy were observed for dexmedetomidine, propofol, and sleep, but changes were larger for propofol compared to dexmedetomidine and sleep. Quantitative comparisons between changes in delta power and network entropy suggest that unresponsiveness under dexmedetomidine produces a similar brain state to that observed during N2 sleep.

**Conclusions:** While delta power, functional connectivity, and network entropy all showed changes during propofol, dexmedetomidine, and sleep, the magnitudes of these changes suggest that dexmedetomidine is more similar than propofol to sleep, specifically to N2 sleep.

## Introduction

Anesthetics can be used experimentally to study both the neural correlates of consciousness and mechanisms underlying its loss and recovery^1–3^. Natural sleep allows for complementary observations without external pharmacological manipulation^4^. Changes in brain activity patterns and connectivity common to unresponsiveness under anesthesia and sleep may be neural correlates of altered or absent consciousness, especially changes between behavioral states qualitatively different in consciousness level.

We have previously shown similarities in resting state (RS) functional connectivity and cortical network organization between NREM sleep and propofol anesthesia using intracranial electroencephalography (iEEG)^5,6^. However, propofol and sleep differ in the spatial pattern and activity/inactivity balance of slow oscillations^7,8^. Propofol is associated with alpha and beta oscillations, but these are not organized into spindles like in sleep^8,9^. In addition, responsiveness to local electrical stimulation was more affected during unarousable propofol anesthesia than sleep^10^.

Dexmedetomidine is an α_2_-adrenergic agonist used clinically as a sedative and as an adjunct to general anesthesia^11,12^. The sedative mechanisms of dexmedetomidine overlap with mechanisms of sleep to produce “arousable sedation”^11–14^, with sleep-like features in the EEG including spindles^15–17^. There is extensive evidence that dexmedetomidine targets neurons in the locus coeruleus and hypothalamus that regulate sleep homeostasis^13,18,19^. However, there is some evidence that electrophysiological activity patterns during dexmedetomidine and sleep are distinct^20,21^ and mixed evidence as to whether dexmedetomidine anesthesia can substitute for sleep^22–24^.

Here, we compare quantitatively the changes observed during dexmedetomidine, propofol, and sleep in spectral power, network entropy, and functional connectivity to see which changes are particular to a certain drug or to natural sleep and which suggest shared mechanisms for altered or loss of consciousness.

## Methods

### Participants

Participants were adult neurosurgical epilepsy patients being monitored intracranially prior to surgical intervention from 2017-2024 without previous resection surgery or major cerebral abnormalities. All participants with data available were used for analysis. Research protocols were approved by the University of Iowa Institutional Review Board (IRB ID# 201804807). Written informed consent was obtained from all participants. Research participation did not interfere with clinical data acquisition, and participants could rescind consent at any time without interrupting clinical evaluation.

Task-free (RS) recordings were obtained from 10 participants (4 female) during induction of anesthesia using dexmedetomidine, 19 participants (11 female) during induction with propofol, and 24 participants (13 female) during natural overnight sleep. Similar analyses have been published previously using data from subsets of the propofol (n=14) and sleep (n=15) participants presented here^5,6^. In the present study, we present recordings during dexmedetomidine anesthesia and use the previously published data and subsequent replication data for comparison.

### Anesthesia recordings

Anesthesia recordings were obtained just prior to electrode removal surgery. Propofol experiments used a gradual infusion (50-150 μg/kg/min) except for four cases where a propofol bolus (180-300 mg) was used at the discretion of the clinical team. Dexmedetomidine experiments used a 0.25 μg/kg bolus followed by 0.2-4 μg/kg/hr, which is higher dosage than routinely used clinically because we were targeting unresponsiveness. To ensure patient safety, the dexmedetomidine infusion was administered at the highest dose for a maximum of 15 minutes, with hemodynamics continuously observed by the anesthesiologist. No participants had cardiac comorbidities. Neither bradycardia, hypotension, nor changes in the electrocardiogram were observed in any participant, and emergence was not delayed. Others have observed no differences in adverse events with higher doses of dexmedetomidine compared to standard doses, even during prolonged administration^25–29^. We note that a higher dose of dexmedetomidine might have been necessary with these patients because some anti-epileptic drugs increase clearance of dexmedetomidine^30^.

RS iEEG data blocks (6-7 minutes) were recorded before (wake anesthesia, “WA”) and during anesthetic infusion^5^. Observer’s Assessment of Alertness/Sedation (OAA/S)^31^ assessments immediately before and after recording blocks were used to label blocks as either sedated (“S”, OAA/S≥3) or unresponsive (“U”, OAA/S≤2)^32^. If behavioral state changed from S to U during a block, the first minute was labeled S, last minute was labeled U, and data between was unused.

### Sleep recordings

Sleep recordings were performed in participants without clinical contraindications who consented to additional scalp electroencephalography and electromyography electrodes needed for polysomnography.

Sleep was staged as wake (WS), non-REM (N1, N2, N3), or REM (R) at 30-second resolution. Sleep recordings occurred median (IQR) 6.3 days (5.6 days, 7.4 days) prior to anesthesia recordings. See Supplementary Methods for more details.

### Region-of-interest assignment

Anatomical reconstructions and assignment of recording contacts to regions of interest (ROIs) was performed as in previous work^5,6^ using co-registered T1-weighted MRI and computed tomography.

### Preprocessing iEEG

Analysis and preprocessing of iEEG was performed using custom software written in MATLAB (v2024a; MathWorks, Natick, MA, USA). Recording sites in seizure foci, in the white matter, or outside the brain were omitted from analysis. The remaining data were subjected to an automated channel and artifact rejection process as described previously^6^. See Supplementary Methods for more details.

### Band power

Time-frequency representations were obtained using the DBT^33^ in the following bands: delta 1-4 Hz, theta 4-8 Hz, alpha 8-14 Hz, beta 14-30 Hz, gamma 30-50 Hz. Power in each band was normalized by bandwidth and averaged across time and frequency within a band in one-minute segments and converted to decibels (dB) relative to 1 μV^2^/Hz.

### Functional connectivity

Pairwise functional connectivity was calculated from DBT time-frequency representations using the debiased weighted phase lag index (wPLI)^34^ and orthogonalized envelope correlations^35^.

wPLI was calculated in the alpha band. Alpha-band wPLI was previously associated with changes in arousal state^5^, and has been commonly used in studies of phase-locked connectivity associated with differences in consciousness^36–39^. As in previous work, we summarized patterns of alpha-band wPLI that change with arousal state in two ways^5^. First, we examined bias in mean wPLI within the “front” versus mean wPLI within the “back” of the brain (see **Table 1** for the list of ROI assignments to Front/Back for this purpose). For this comparison, we omitted two participants whose only “front” coverage was in Amyg or Hipp, with no neocortical coverage in the “front”. Second, we examined mean connectivity specifically in “long-range” wPLI, defined as the top 50% longest physical distances between recording sites among all pairwise distances.

**Table 1.**
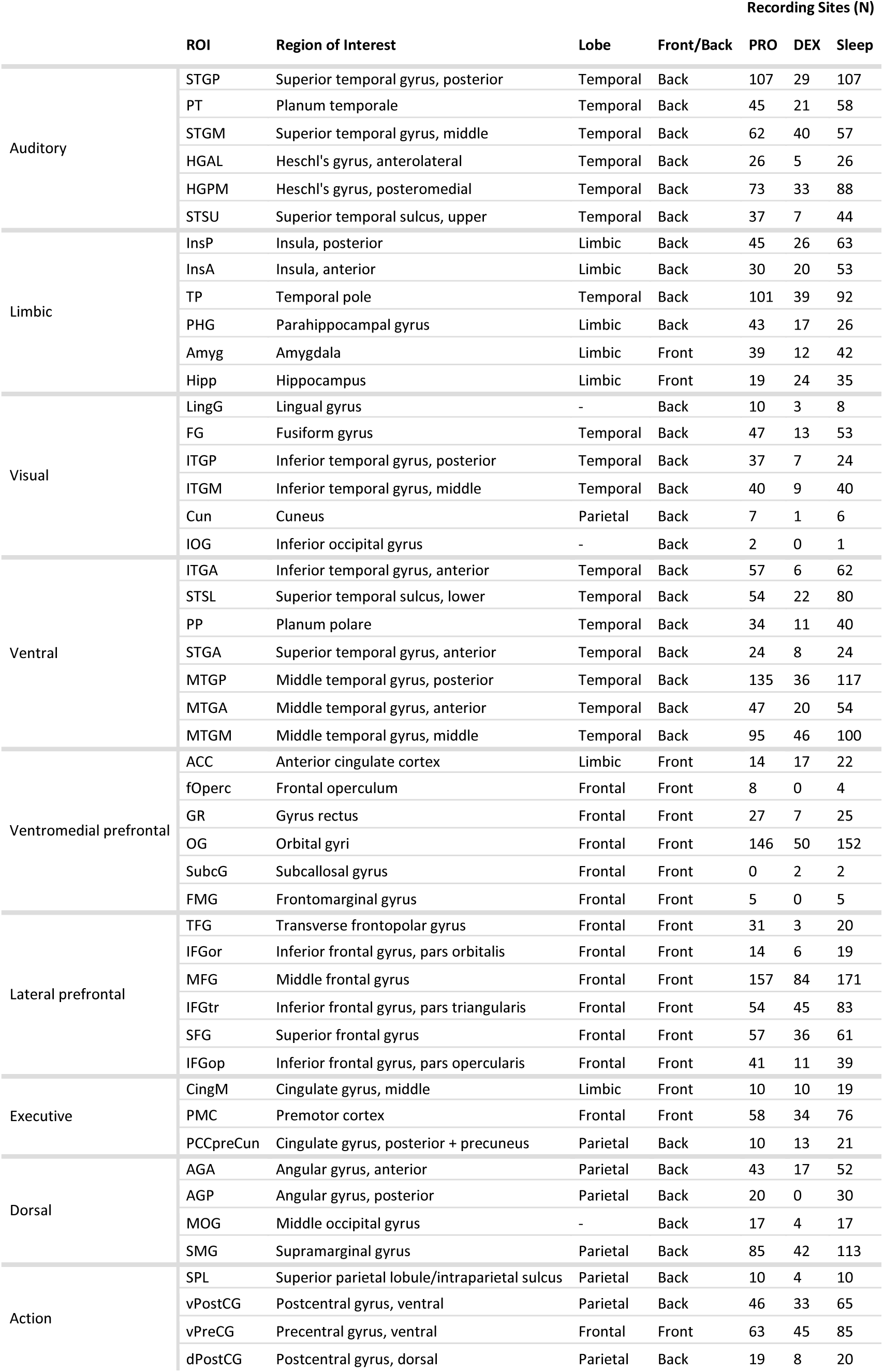
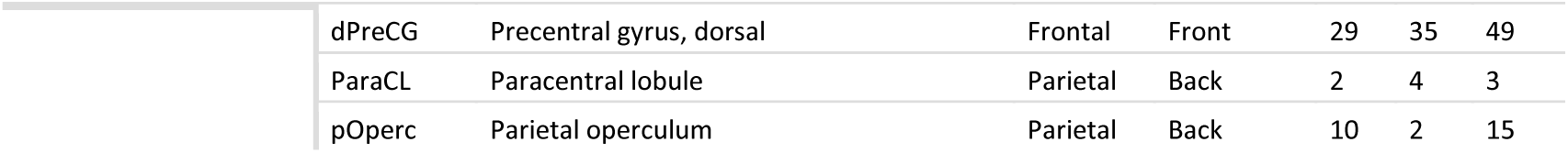
Number of recording sites by ROI and experiment. Lobe refers to lobes used for analysis of band power. Front/back refers to groupings used for wPLI analysis.

|  |  |  |  |  | Recording Sites (N) |  |  |
| --- | --- | --- | --- | --- | --- | --- | --- |
|  | ROI | Region of Interest | Lobe | Front/Back | PRO | DEX | Sleep |
| Auditory | STGP | Superior temporal gyrus, posterior | Temporal | Back | 107 | 29 | 107 |
|  | PT | Planum temporale | Temporal | Back | 45 | 21 | 58 |
|  | STGM | Superior temporal gyrus, middle | Temporal | Back | 62 | 40 | 57 |
|  | HGAL | Heschl's gyrus, anterolateral | Temporal | Back | 26 | 5 | 26 |
|  | HGPM | Heschl's gyrus, posteromedial | Temporal | Back | 73 | 33 | 88 |
|  | STSU | Superior temporal sulcus, upper | Temporal | Back | 37 | 7 | 44 |
| Limbic | InsP | Insula, posterior | Limbic | Back | 45 | 26 | 63 |
|  | InsA | Insula, anterior | Limbic | Back | 30 | 20 | 53 |
|  | TP | Temporal pole | Temporal | Back | 101 | 39 | 92 |
|  | PHG | Parahippocampal gyrus | Limbic | Back | 43 | 17 | 26 |
|  | Amyg | Amygdala | Limbic | Front | 39 | 12 | 42 |
|  | Hipp | Hippocampus | Limbic | Front | 19 | 24 | 35 |
| Visual | LingG | Lingual gyrus | - | Back | 10 | 3 | 8 |
|  | FG | Fusiform gyrus | Temporal | Back | 47 | 13 | 53 |
|  | ITGP | Inferior temporal gyrus, posterior | Temporal | Back | 37 | 7 | 24 |
|  | ITGM | Inferior temporal gyrus, middle | Temporal | Back | 40 | 9 | 40 |
|  | Cun | Cuneus | Parietal | Back | 7 | 1 | 6 |
|  | IOG | Inferior occipital gyrus | - | Back | 2 | 0 | 1 |
| Ventral | ITGA | Inferior temporal gyrus, anterior | Temporal | Back | 57 | 6 | 62 |
|  | STSL | Superior temporal sulcus, lower | Temporal | Back | 54 | 22 | 80 |
|  | PP | Planum polare | Temporal | Back | 34 | 11 | 40 |
|  | STGA | Superior temporal gyrus, anterior | Temporal | Back | 24 | 8 | 24 |
|  | MTGP | Middle temporal gyrus, posterior | Temporal | Back | 135 | 36 | 117 |
|  | MTGA | Middle temporal gyrus, anterior | Temporal | Back | 47 | 20 | 54 |
|  | MTGM | Middle temporal gyrus, middle | Temporal | Back | 95 | 46 | 100 |
| Ventromedial prefrontal | ACC | Anterior cingulate cortex | Limbic | Front | 14 | 17 | 22 |
|  | fOperc | Frontal operculum | Frontal | Front | 8 | 0 | 4 |
|  | GR | Gyrus rectus | Frontal | Front | 27 | 7 | 25 |
|  | OG | Orbital gyri | Frontal | Front | 146 | 50 | 152 |
|  | SubcG | Subcallosal gyrus | Frontal | Front | 0 | 2 | 2 |
|  | FMG | Frontomarginal gyrus | Frontal | Front | 5 | 0 | 5 |
| Lateral prefrontal | TFG | Transverse frontopolar gyrus | Frontal | Front | 31 | 3 | 20 |
|  | IFGor | Inferior frontal gyrus, pars orbitalis | Frontal | Front | 14 | 6 | 19 |
|  | MFG | Middle frontal gyrus | Frontal | Front | 157 | 84 | 171 |
|  | IFGtr | Inferior frontal gyrus, pars triangularis | Frontal | Front | 54 | 45 | 83 |
|  | SFG | Superior frontal gyrus | Frontal | Front | 57 | 36 | 61 |
|  | IFGop | Inferior frontal gyrus, pars opercularis | Frontal | Front | 41 | 11 | 39 |
| Executive | CingM | Cingulate gyrus, middle | Limbic | Front | 10 | 10 | 19 |
|  | PMC | Premotor cortex | Frontal | Front | 58 | 34 | 76 |
|  | PCCpreCun | Cingulate gyrus, posterior + precuneus | Parietal | Back | 10 | 13 | 21 |
| Dorsal | AGA | Angular gyrus, anterior | Parietal | Back | 43 | 17 | 52 |
|  | AGP | Angular gyrus, posterior | Parietal | Back | 20 | 0 | 30 |
|  | MOG | Middle occipital gyrus | - | Back | 17 | 4 | 17 |
|  | SMG | Supramarginal gyrus | Parietal | Back | 85 | 42 | 113 |
| Action | SPL | Superior parietal lobule/intraparietal sulcus | Parietal | Back | 10 | 4 | 10 |
|  | vPostCG | Postcentral gyrus, ventral | Parietal | Back | 46 | 33 | 65 |
|  | vPreCG | Precentral gyrus, ventral | Frontal | Front | 63 | 45 | 85 |
|  | dPostCG | Postcentral gyrus, dorsal | Parietal | Back | 19 | 8 | 20 |

|  |  |  |  |  |  |  |
| --- | --- | --- | --- | --- | --- | --- |
| dPreCG | Precentral gyrus, dorsal | Frontal | Front | 29 | 35 | 49 |
| ParaCL | Paracentral lobule | Parietal | Back | 2 | 4 | 3 |
| pOperc | Parietal operculum | Parietal | Back | 10 | 2 | 15 |

Orthogonalized envelope correlations were calculated in the gamma band, as previously used in an analysis of sleep and propofol anesthesia^6^. Gamma band activity correlates with neuronal firing rates and gamma correlations with blood oxygen-level dependent functional connectivity^40,41^.

### Diffusion map embedding and network entropy

Diffusion map embedding was used to characterize the functional geometry of neural activity as measured by gamma envelope correlations^42,43^. See Supplementary Methods for more details.

The eigenvalue spectrum |λ| of the diffusion map embedding represents the structural dimensionality or complexity of the underlying network that generates the data^44^. To characterize the “structure” versus “flatness” of the eigenvalue spectrum, we use the normalized Shannon entropy of the normalized eigenvalues [λ ^‘^… λ ^‘^]^45^, where:

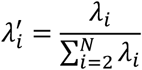

and with **n = N – 1** eigenvalues,

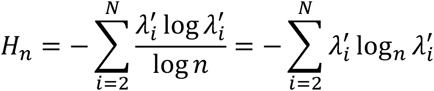

We refer to this normalized entropy H_n_ as “network entropy”.

### Statistical modeling

For each outcome measure, linear mixed effects models were fit with fixed effects for behavioral state (W_DEX_, S_DEX_, U_DEX_, W_PRO_, S_PRO_, U_PRO_, WS, N1, N2, N3, R) and random effects for participant and experiment day using the R^46^ package lme4^47^. In Wilkinson notation the model is:

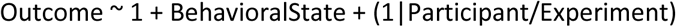

Marginal means for each behavioral state and contrasts between states were estimated from fitted models using the package emmeans^48^. Satterthwaite’s approximation was used for degrees of freedom. Adjustments for multiple comparisons within each model and experiment were performed by simultaneous inference among contrasts using a multivariate t distribution (‘mvt’ option in emmeans). For anesthesia experiments, all three pairwise contrasts were tested. For sleep, planned pairwise contrasts for testing were those between WS and each other state, sequential comparisons between N1 and N2, N2 and N3, and between R and N2, R and N3.

No *a priori* sample size calculations were performed for this work.

### Data and code availability

Data and code necessary to reproduce figures and statistical analyses performed for this manuscript are available at (https://doi.org/10.5281/zenodo.15497531). Raw patient data are available for approved scientific projects with a data transfer and use agreement through The University of Iowa Division of Sponsored Projects at subject to the terms under which participants’ consent was given.

## Results

### Summary

Dexmedetomidine (DEX) experiments were performed in 10 participants (6 also with sleep). Propofol (PRO) experiments were performed in 19 participants (13 also with sleep). Sleep data was used from 5 additional participants who did not have recordings during anesthesia. There were a total of 2,192 recording sites in propofol experiments, 967 in dexmedetomidine, and 2,478 in sleep (**Table 1**, **Figure 1**). A total of 12,313 minute-long segments of data were analyzed (**Supplementary Table 1)**.

**Figure 1.**
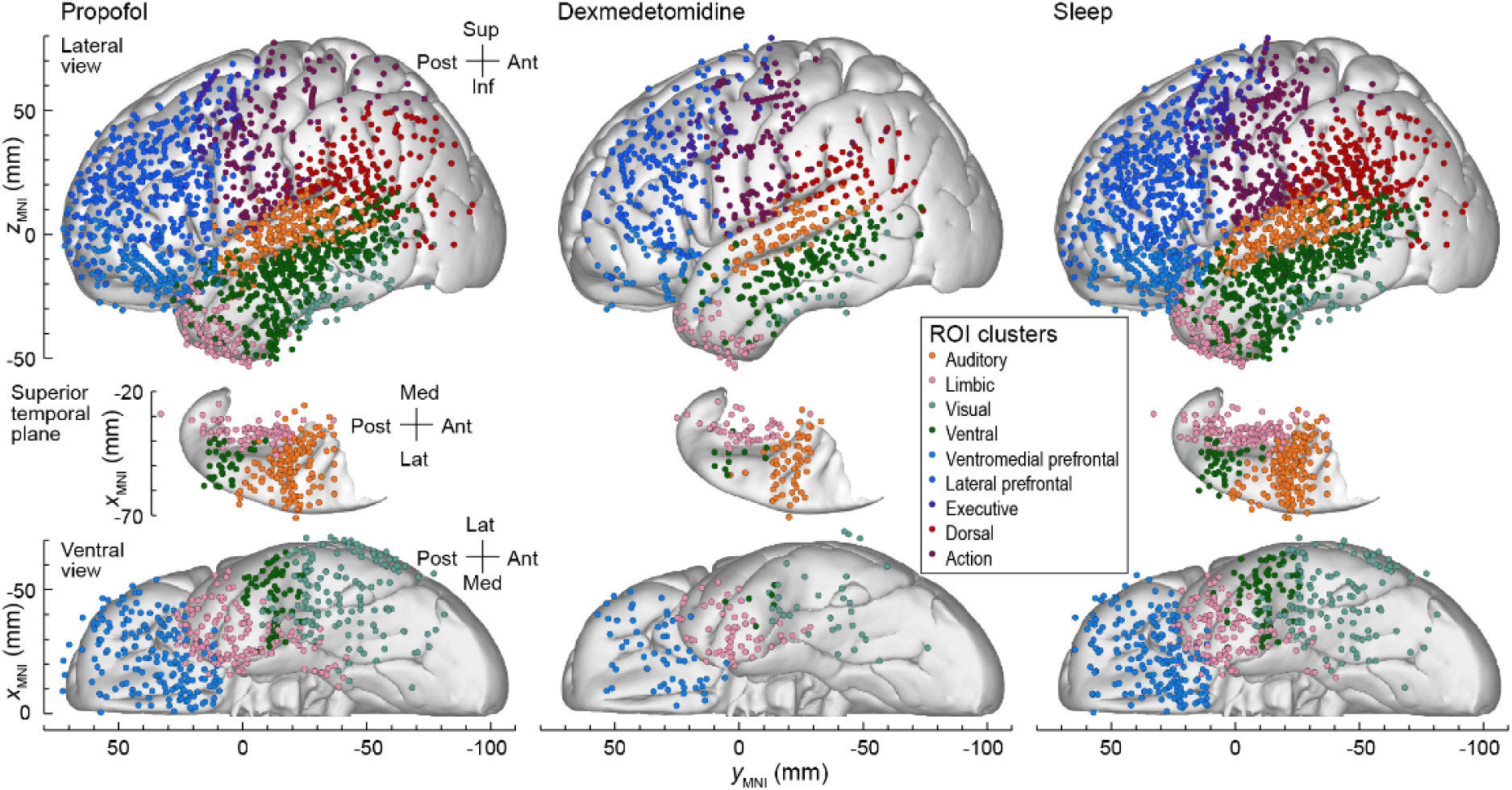
Recording site coverage for all experiments. All recording sites are reflected to the left hemisphere for display. Colors group recording sites into regional (ROI) clusters that are derived from previous work^43^.

### Band power

As expected^49,50^ and consistent with our previous work^5,6^, differences in behavioral state (i.e., stages of sleep and anesthesia) associated with changes in the iEEG power spectrum (**Figure 2a**). Increases in delta power were observed for both anesthetic agents and sleep.

**Figure 2.**
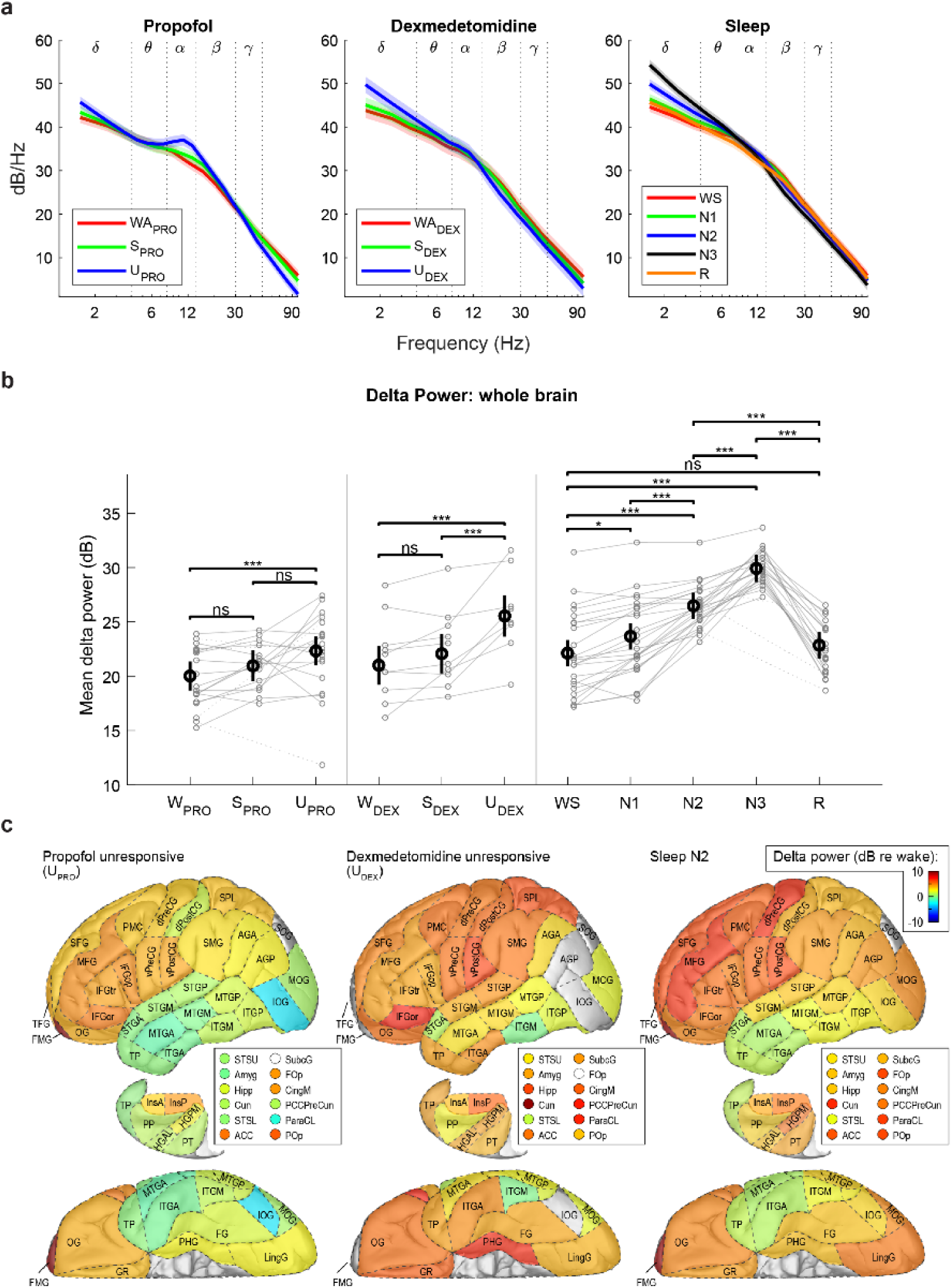
Effect of behavioral state on the frequency content of resting state brain activity. **a)** Average power spectra by behavioral state. Solid lines are predicted means and shaded areas depict 95% confidence intervals for the mean from a linear mixed effects model. **b)** Summary of delta power changes. Gray symbols with lines are means of log delta power across recording sites for each participant; dashed lines are used when no data for an intervening behavioral state were available for that participant. Black symbols and error bars are predicted marginal means and 95% confidence intervals for the mean across participants. * *p* < 0.05, *** *p* < 0.001, ns = not significant. **c)** Delta power changes while unresponsive under anesthesia or in N2 sleep, relative to wake. Colored ROIs show predicted mean differences across recording site and participant for each ROI, in units of dB relative to W_PRO_, W_DEX_, or WS. Top: lateral view, middle: superior temporal plane, bottom: ventral view. ROIs that are not visible in one of the three views are displayed separately as labeled symbols. ROIs without recording sites for the comparison depicted are shaded gray.

Estimated mean delta power across participants and single-participant means across all recording sites are presented in **Figure 2b**. Comparisons between groups are summarized in **Table 2**. As expected, delta power increased with increasing depth of dexmedetomidine and propofol anesthesia (W < S < U) and also increased with progressively deeper states of non-REM sleep (WS < N1 < N2 < N3); R was not significantly different from WS.

**Table 2.**
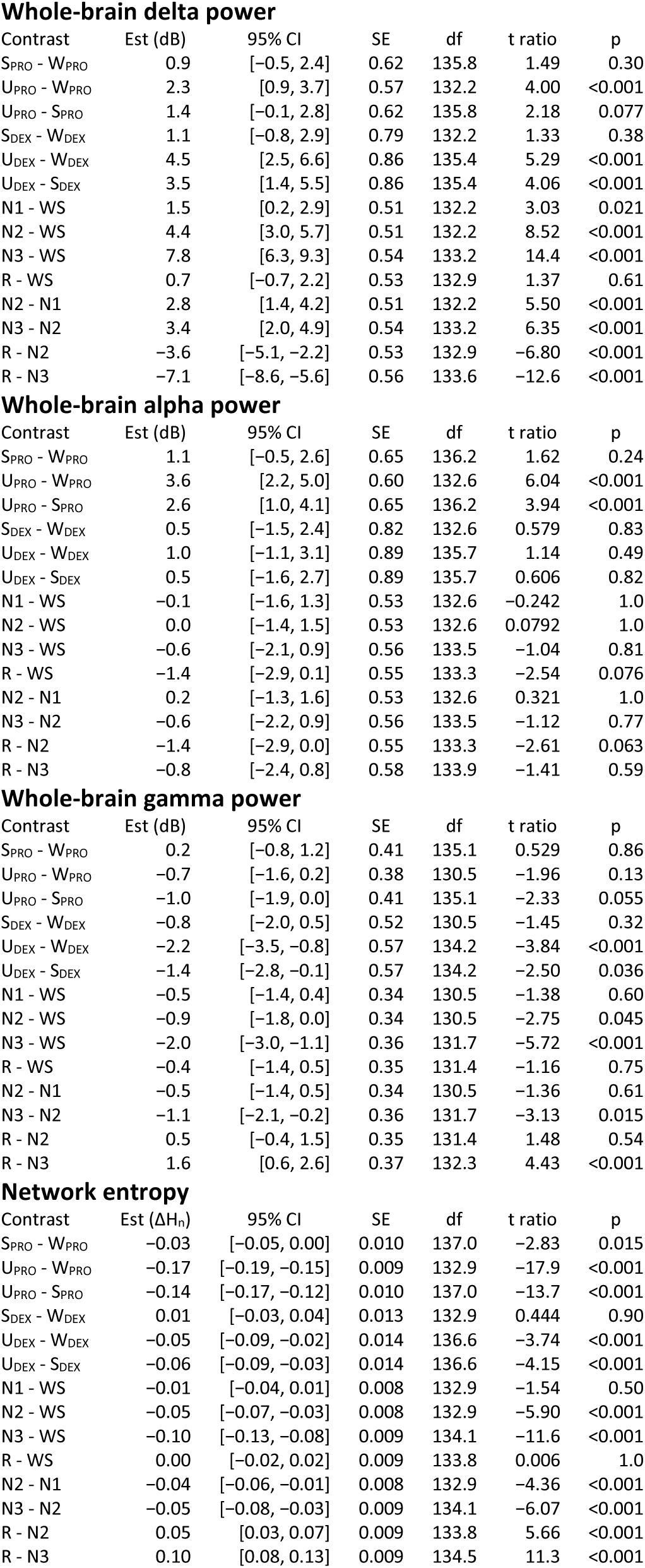
Planned contrasts for whole-brain power outcomes and network entropy. Est = estimated difference. CI = confidence interval. SE = standard error. df = degrees of freedom using Satterthwaite approximation.

Regional changes in delta power are shown in **Figure 2c**. To quantify, we estimated mean delta power in frontal, parietal, temporal, and limbic lobes for each state (**Supplementary Figure 1)** and compared between states (**Supplementary Table 2**). Significant delta power increases in U_DEX_ relative to W_DEX_ and in N2 relative to WS were observed in each lobe. (N2 was used for comparison because fewer segments were staged as N3 and some participants had no N3 sleep.) However, while delta power increases were also seen in U_PRO_ for frontal, parietal, and limbic lobes, in temporal lobe the change was not significantly different from zero.

For other frequency bands, power changes with depth of anesthesia were not consistent across dexmedetomidine, propofol, and sleep (**Supplementary Figure 2; Supplementary Table 3**). Alpha power increased significantly in U_PRO_ compared to S_PRO_ and W_PRO_, but for dexmedetomidine alpha power did not change significantly with behavioral state, nor were there significant differences between WS and non-REM sleep. Frontal alpha specifically showed much larger differences between states for propofol (for example, U_PRO_ – S_PRO_ = 6.0, 95% CI [4.0, 8.1] for frontal alpha, versus U_PRO_ – S_PRO_ = 2.6, 95% CI [1.0, 4.1] for the whole brain), but differences between U_DEX_ and S_DEX_ or between wake and non-REM sleep were not significant (**Supplementary Table 4**).

In contrast, gamma power decreased significantly in U_DEX_ compared to S_DEX_ and W_DEX_, as well as from WS to N2 and N3, but gamma power in U_PRO_ was not significantly different from W_PRO_ (**Supplementary Figure 2; Supplementary Table 3**).

### Gamma envelope network entropy

We previously showed that network complexity derived from RS functional connectivity changed consistently during sleep and propofol anesthesia, suggesting common underlying mechanisms of loss of consciousness^6^. Here, we determined whether the same was true for dexmedetomidine anesthesia. We treated functional connectivity matrices (calculated using pairwise correlations of gamma power envelopes) as graphs that can be traversed by random walk, and characterized these graphs using diffusion map embedding^6,43,51^. When functional connectivity matrices are sorted, a “blocked” structure is visually apparent: groups of recording sites share stronger connectivity with some groups but not others

(**Figure 3)**. More organized diffusion matrices have less entropy in their eigenvalue spectrum (H_n_) and the network can be described at the same fidelity using fewer dimensions^6^. We observed similar changes in H_n_ with dexmedetomidine anesthesia as we did with propofol and sleep (**Figure 3c**; Error! Reference source not found.). H_n_ decreased with increasing depth of propofol and dexmedetomidine anesthesia (W > S > U). H_n_ also decreased with increasing depth of NREM sleep (WS > N1 > N2 > N3), but R was not significantly different from WS. For direct comparison to previous work, we also quantify the eigenvalue spectrum using D_E_ (**Supplementary Figure 3; Supplementary Table 5**).

**Figure 3.**
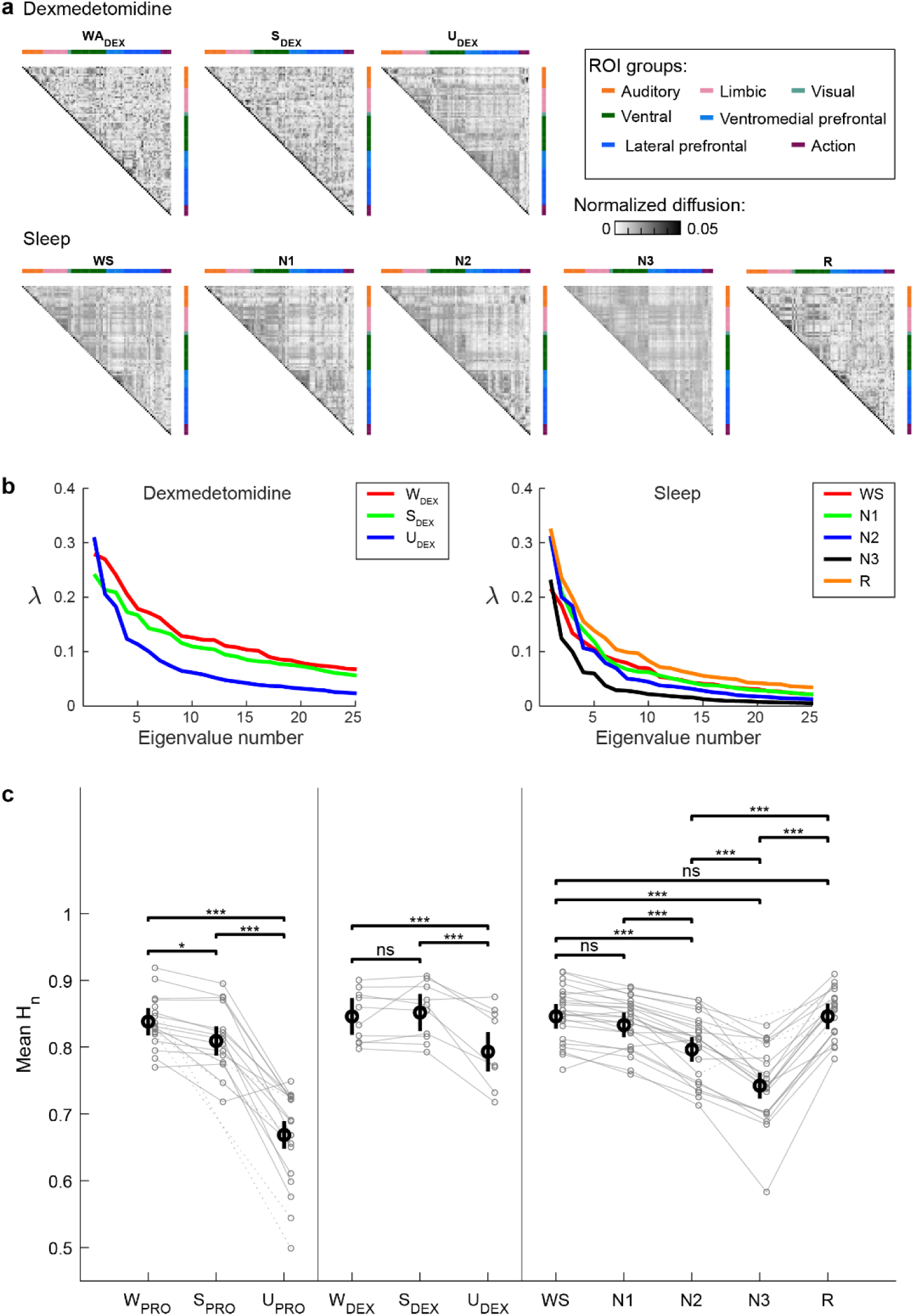
Network entropy of gamma envelope correlations changes with dexmedetomidine anesthesia and sleep. **a)** Example gamma envelope diffusion matrices from one participant. Each matrix is derived from one minute of data. **b)** Eigenvalue spectra calculated from the diffusion matrices in **(a)**. Network entropy is derived from these spectra; for these examples, H_n_(W_DEX_) = 0.87, H_n_(S_DEX_) = 0.86, H_n_(U_DEX_) = 0.77, H_n_(WS) = 0.79, H_n_(N1) = 0.76, H_n_(N2) = 0.73, H_n_(N3) = 0.69, and H_n_(R) = 0.81. **c)** Summary of network entropy across experiments and participants. Gray symbols with lines are means across segments for each participant; dashed lines are used when no data for a given behavioral state were available for that participant. Black symbols and error bars are predicted marginal means and 95% confidence intervals for the mean across participants. * *p* < 0.05, *** *p* < 0.001, ns = not significant.

### Alpha wPLI

In a previous study on five of the participants used in this study, certain patterns in alpha wPLI appeared to change consistently with propofol anesthesia and sleep^5^. Specifically, for both propofol unresponsiveness and deep sleep, 1) alpha wPLI increased within the “front” of the brain relative to the “back” (see Methods), and 2) increased between regions that were physically distant (“long-range”).

In the present expanded dataset, we also found significant increases in front versus back wPLI when participants were unresponsive (U) under propofol or dexmedetomidine anesthesia compared to W or S, and for nREM sleep compared to WS or R (**Supplementary Figure 4a; Supplementary Table 6**). Long-range alpha wPLI also increased in unresponsive (U compared to W or S in both propofol and dexmedetomidine anesthesia (**Supplementary Figure 4b; Supplementary Table 6**). However, we did not find significant changes in long-range connectivity from WS to non-REM sleep.

### Dexmedetomidine parallels sleep in delta power and network entropy

Although there are commonalities between sleep and both anesthetics in the direction of change across several measures (power, network entropy, wPLI), the relative magnitudes of these changes were not the same. In particular, in the two-dimensional space defined by network entropy and delta power, the data for dexmedetomidine overlies sleep, while the data for propofol diverge markedly (**Figure 4**). When delta power increased with sleep or dexmedetomidine, network entropy decreased by a corresponding amount, such that N2 sleep and U_DEX_ are similar in both measures. In contrast, large decreases in network entropy for propofol were accompanied by proportionally smaller changes in delta power. These results support the idea that the effects of dexmedetomidine on brain activity and connectivity are more “sleep-like” compared to the distinct changes with propofol.

**Figure 4.**
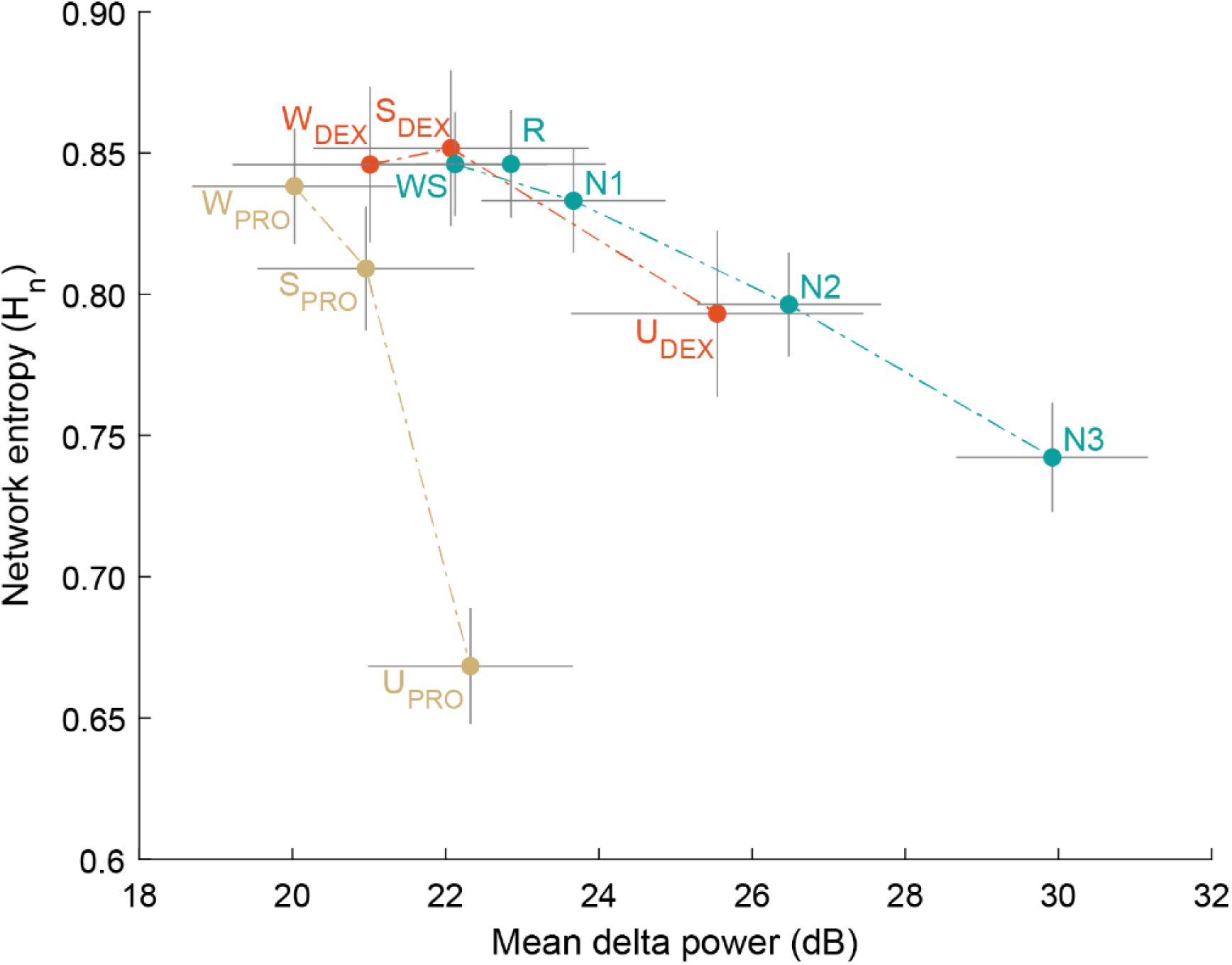
Relative changes in delta power and network entropy. Labeled points are predicted means; horizontal and vertical lines are 95% confidence intervals for delta power and network entropy, respectively. Dot-dashed lines connect points in order of deeper anesthesia or stages of non-REM sleep.

## Discussion

### Summary

Changes in consciousness result from a variety of pharmacological, pathophysiological, and natural conditions. Comparisons of brain activity and connectivity across these conditions can produce insight into neural correlates of consciousness and mechanisms of loss and recovery of consciousness^5,6,21,52–54^. Here, we show that dexmedetomidine anesthesia is accompanied by changes in brain activity and connectivity that generally align with those observed previously for propofol anesthesia and overnight sleep, and are most consistent with the latter.

Delta power increased with depth of anesthesia and with progressively deeper NREM sleep. Delta power has often been suggested to have predictive power for loss and recovery of consciousness during anesthesia^9,50^ as is the case for sleep^4,55^. Slow waves, which contribute importantly to the delta band signal, are prominent during both anesthesia and NREM sleep and interfere with cognition by disrupting information sharing within and between brain regions^56^. However, delta power can also be elevated without loss of consciousness^57–59^. Here, we show that delta power changes during unresponsiveness under dexmedetomidine anesthesia were similar to those observed during N2 sleep, whereas changes with propofol were smaller and more localized.

Network entropy (H_n_), measured from the eigenvalue spectrum of RS functional connectivity, decreased with anesthesia depth and depth of NREM sleep. Changes in network entropy capture changes in functional integration and differentiation of brain signals^6^, aligning these results with the growing body of work showing that during anesthesia and sleep, brain networks shift the balance of integration and segregation toward the latter^52,60,61^. Connectivity within the front versus back of the brain also showed consistent changes for dexmedetomidine, propofol, and natural sleep.

### Quantitative differences among dexmedetomidine, propofol, and sleep

An important contribution of the current study is the observation that dexmedetomidine produces changes in brain activity and network structure that are more comparable to NREM sleep than are those observed under propofol anesthesia. This observation stems from moving beyond hypothesis testing to a quantitative comparison of effect sizes.

Significance thresholds based on p-values contain little information about effect magnitude; “significant” changes can associate with either small or large effects depending on sample size, study design, and randomness. Researchers are encouraged to instead use measures of “effect size”^62^, but interpreting effect sizes is difficult with outcomes that lack a tangible or familiar understanding. By comparing dexmedetomidine and propofol to each other and to sleep, we introduce internal reference points to interpret the magnitude of differences. For example, the delta power increase is of similar magnitude for dexmedetomidine and sleep but only half as large for propofol in a comparable behavioral state. The magnitude of changes in H_n_ corresponded better to qualitative perceptions about the different behavioral states than delta power, such as propofol unresponsiveness being a less-arousable state than sleep or dexmedetomidine anesthesia.

### Caveats

This study sampled only from participants with a neurological disorder and may not generalize to a broader population^5,6^. Electrodes were placed based on clinical needs rather than research goals, contributing to variability across participants. History of epilepsy, use of medications including antiseizure medications, and the hospital environment can all affect sleep and anesthesia sensitivity^63–67^. We mitigated these confounding influences by conditioning on response to command rather than dose/concentration of drugs and by using polysomnography to identify behavioral states in sleep experiments. Not all participants in our cohort slept deeply in the hospital, therefore we focused comparisons on N2 rather than N3 sleep. Despite differing clinical characteristics and differing medication across participants, we found common differences in neurophysiology between behavioral states in anesthesia and sleep.

Dreaming is common under dexmedetomidine and propofol anesthesia^20,68–71^, as it is during N1 and N2 sleep^72^. Dreaming is considered a form of consciousness accompanied by a state of disconnection^21,73^. We cannot know for certain whether unresponsive participants were unconscious or dreaming, but we can relate our results to the probability of (un)consciousness from serial awakening experiments, with the lowest probability in N3 sleep and a high probability of dreaming consciousness in REM^72^. For example, H_n_ was lowest in N3 sleep and similar to wake in REM, suggesting that H_n_ indexes consciousness as opposed to responsiveness.

## Conclusion

Previous evidence for the sleep-like characteristics of dexmedetomidine anesthesia include shared molecular targets with natural sleep^13,14,24^, sleep-like features in the EEG like spindles^15–17^, and clinical impressions of sedated patients^11,12,74^. In our analyses, unresponsiveness under dexmedetomidine anesthesia is most comparable to N2 sleep in delta band power and H_n_ and distinct from unresponsiveness under propofol anesthesia.

## Author Contributions

Study conception and design: BK, KN, MB

Data collection and curation: BK, ED, RM, HK, KN Data analysis: BK, MB

Data interpretation: BK, RS, KN, MB Manuscript draft: BK, MB

Manuscript review and editing: BK, ED, RM, RS, KN, MB

Reviewed and approved the final manuscript submitted: all authors

## Acknowledgements

The authors are grateful to Haiming Chen, Ariane Rhone, Christopher Garcia, and Christopher Kovach for assistance with data collection and analysis.

## Supplementary Methods

### Sleep recordings

For participants with multiple nights of polysomnography, we chose a single night to analyze, first prioritizing the night with the most different sleep stages available (because N3 and REM were not observed every night), and second prioritizing the night with the most total time scored as N1, N3, and REM, since these stages were typically less common. One participant (403L) experienced multiple seizures overnight so we only analyzed the first half of the night, before seizures.

### iEEG artifact and channel rejection

Initial denoising including removal of 60 Hz electrical noise was performed using the demodulated band transform (DBT)^33^. Channels were excluded if they had unusually high or low power (±3.5 SD relative to all channels) in any frequency band. Voltage deflections exceeding 10 SD in any channel, high-gamma power exceeding 5 SD, or segments with saturation of the amplifier (“clipping”) were marked as artifacts. If particular channels had unusually frequent artifacts or amplifier saturation, they were excluded entirely, otherwise artifact exclusions applied across all channels and for 100 ms before and after each event. Data were divided into 1-minute segments for analysis; segments with more than 20 seconds of data marked as artifact were omitted entirely. A spatial filter derived from singular value decomposition of the high-pass (200 Hz) filtered data was applied to the broadband signal to remove common zero-lag signal shared from the recording environment as previously^6^.

### Diffusion map embedding details

The similarity matrix **K =** [*k*(*i*,*j*)] is obtained from the cosine similarity of vectors *i,j* in the functional connectivity adjacency matrix (with the diagonal defined as equal to 1), then normalized by the diagonal degree matrix **D** to yield a diffusion matrix **P = D**^-1^**K** that represents a random walk on the graph. The eigenvalues [λ_2_… λ_N_] of P represent the variance along each embedding dimension. The first eigenvalue λ_1_ is 1 because P is a stochastic matrix, with an associated steady-state eigenvector that is made constant through the normalization by degree; we focus on the other interesting dimensions.

The eigenvalue spectrum |λ| (also called the “spectrum”; we use “eigenvalue spectrum” to avoid confusion with other associations of “spectrum”) represents the structural dimensionality or complexity of the underlying network that generates the data^44^. Unstructured or “random” connectivity will have a flat eigenvalue spectrum with equal eigenvalue magnitudes; if connectivity is highly structured and depends on a small number of features, those features will be mapped to dimensions having large eigenvalues and the eigenvalues for other dimensions will be small.

To characterize the “structure” versus “flatness” of the eigenvalue spectrum, we use the normalized Shannon entropy of the normalized eigenvalues [λ ^‘^… λ ^‘^]^45^, where:

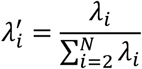

and with **n = N – 1** eigenvalues,

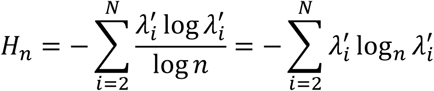

We refer to this normalized entropy H_n_ as “network entropy”.

We have previously used the perplexity (exponentiation) of quadratic spectral entropy divided by the total dimensionality (the “normalized effective dimensionality”); we include this measure for comparison with that previous work^6^:

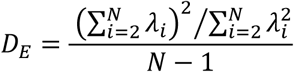

Both quantifications involve calculations of entropy on the eigenvalue spectrum, but with different normalizations. For network entropy, we used Shannon entropy corresponding to the Rényi entropy in the limit α → 1; quadratic entropy corresponds to α = 2. Larger values of α weight larger eigenvalues more heavily. We calculated effective dimensionality as the perplexity of the quadratic entropy, then divided by the original (maximum) number of dimensions to obtain a number comparable across participants differing in number of recording sites. Quadratic entropy may be more useful if the goal is to reduce to some conservative number of dimensions, though Shannon entropy can also produce a valid estimate of the effective dimensionality^75^. Here we chose to use the normalized Shannon entropy rather than effective dimensionality because we observe less variance across participants but are still able to distinguish among states. This method also avoids the step of converting entropy to perplexity, which produces values usefully interpretable as the number of dimensions, but which then requires further normalization to compare across participants who differ in maximum dimensionality because of differing numbers of recording sites.

## Supplementary Figures

**Supplementary Figure 1.**
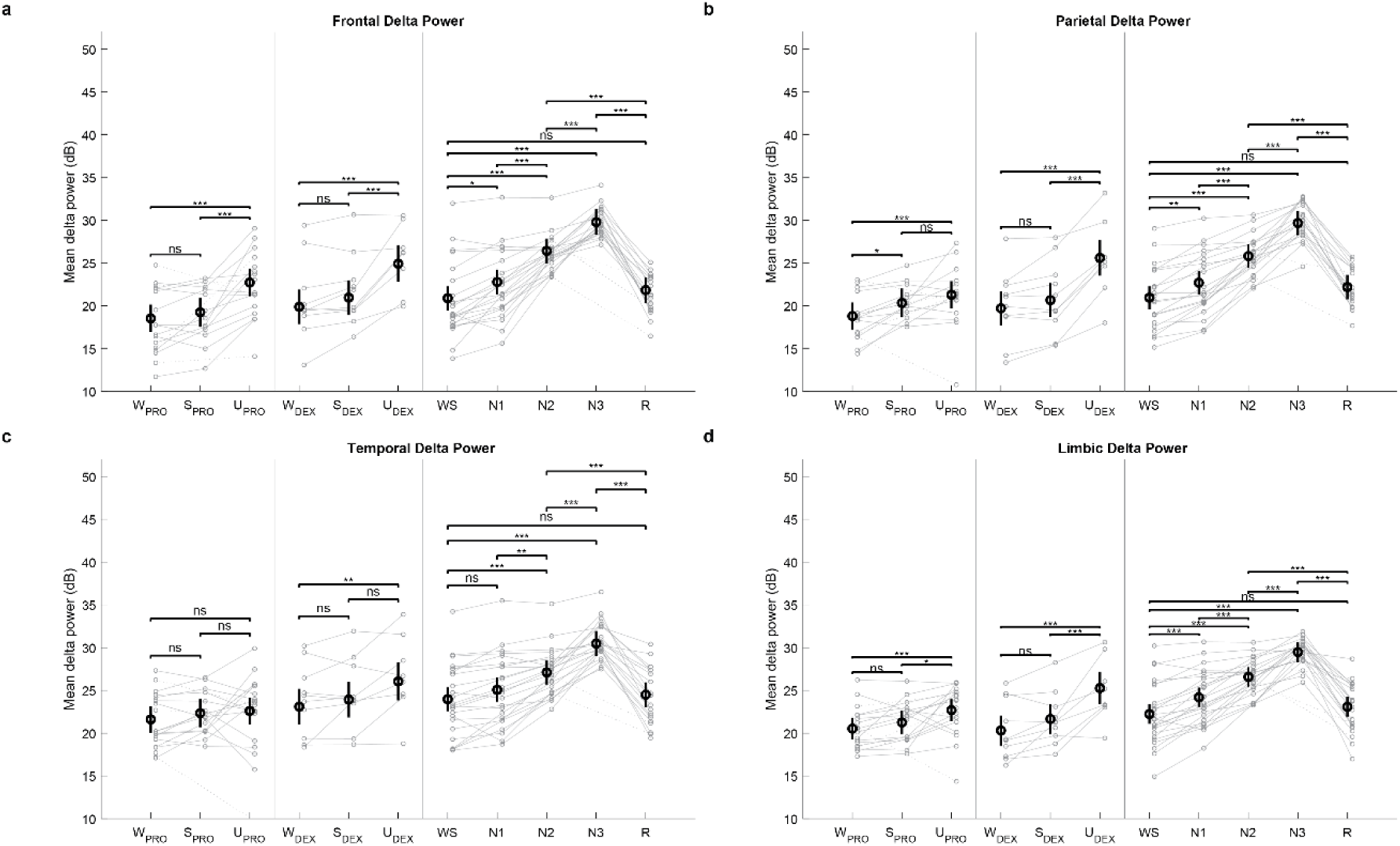
Mean delta power across electrode contacts for each region. **a)** frontal, **b)** parietal, **c)** temporal, **d)** limbic. Units are dB relative to 1 μV^2^/Hz. Gray symbols with lines are means across recording sites for each participant; dashed lines are used when no data for an intervening behavioral state were available for that participant. Black symbols and error bars are predicted marginal means and 95% confidence intervals for the mean across participants. * *p* < 0.05, ** *p* < 0.01, *** *p* < 0.001, ns = not significant.

**Supplementary Figure 2.**
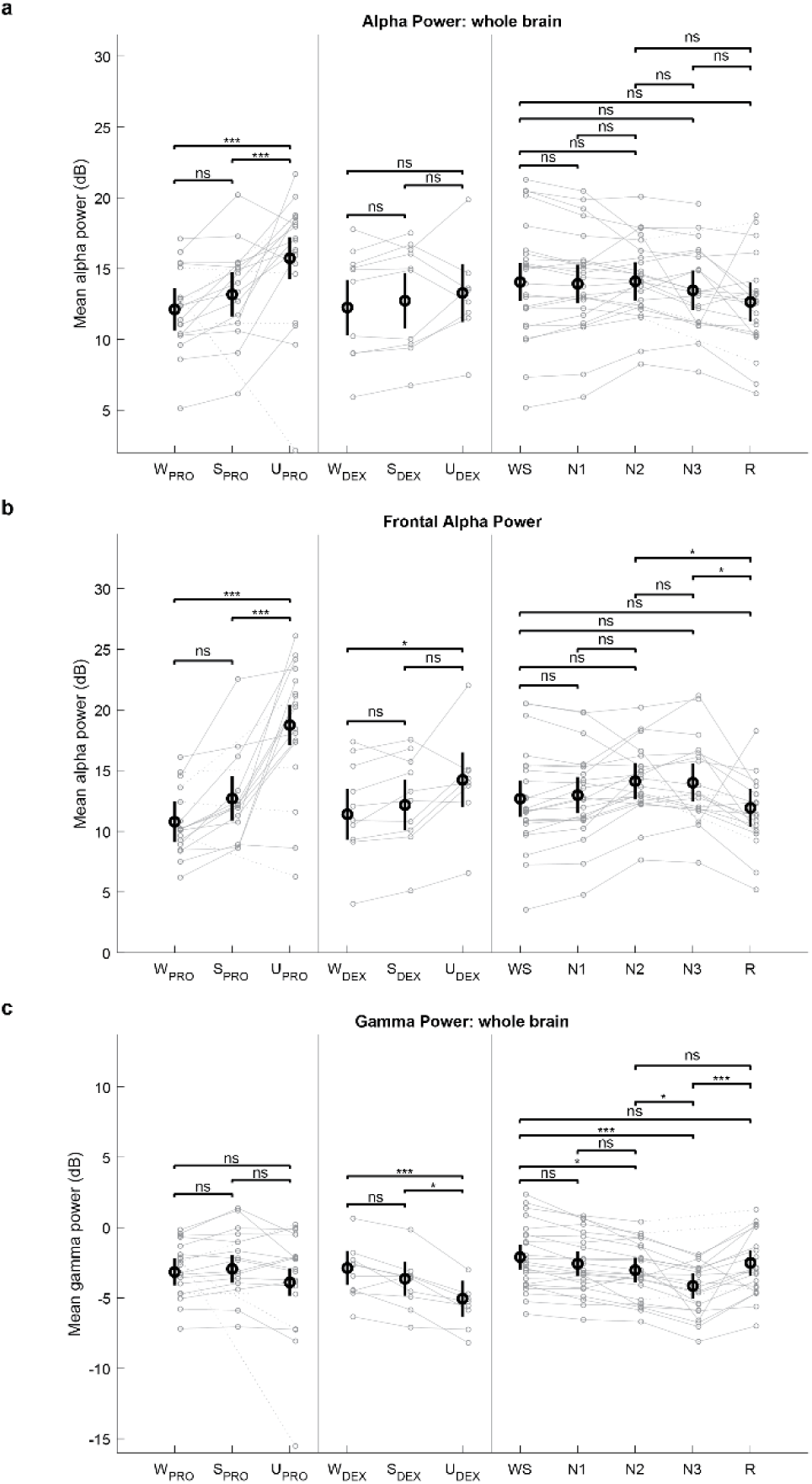
Power in alpha and gamma bands. **a)** Mean alpha power across all recording sites, in dB relative to 1 μV^2^/Hz. Gray symbols with lines are means across recording sites for each participant; dashed lines are used when no data for an intervening behavioral state were available for that participant. Black symbols and error bars are predicted marginal means and 95% confidence intervals for the mean across participants. * *p* < 0.05, ** *p* < 0.01, *** *p* < 0.001, ns = not significant. **b)** Same as **(a)**, for alpha power in the frontal lobe only. **c)** Same as **(a)**, for the gamma band.

**Supplementary Figure 3.**
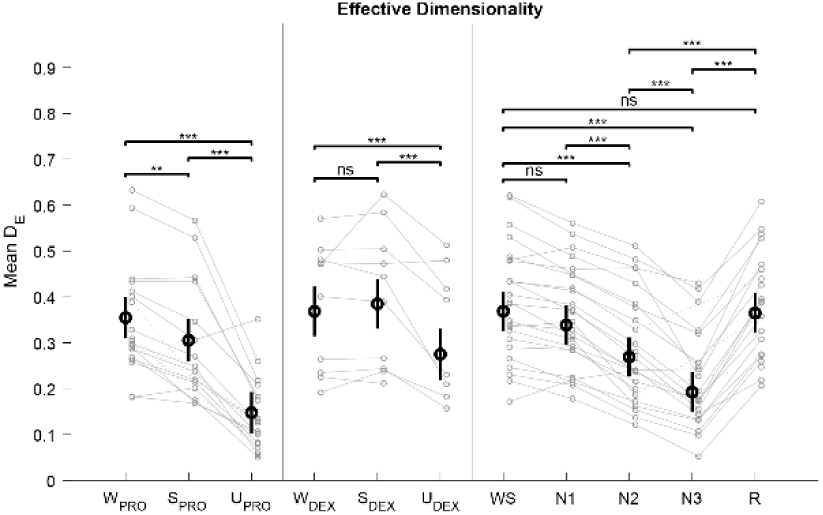
Effective dimensionality decreases with sleep and anesthesia. Gray symbols with lines are means across segments for each participant; dashed lines are used when no data for a given behavioral state were available for that participant. Black symbols and error bars are predicted marginal means and 95% confidence intervals for the mean across participants. * *p* < 0.05, *** *p* < 0.001, ns = not significant

**Supplementary Figure 4.**
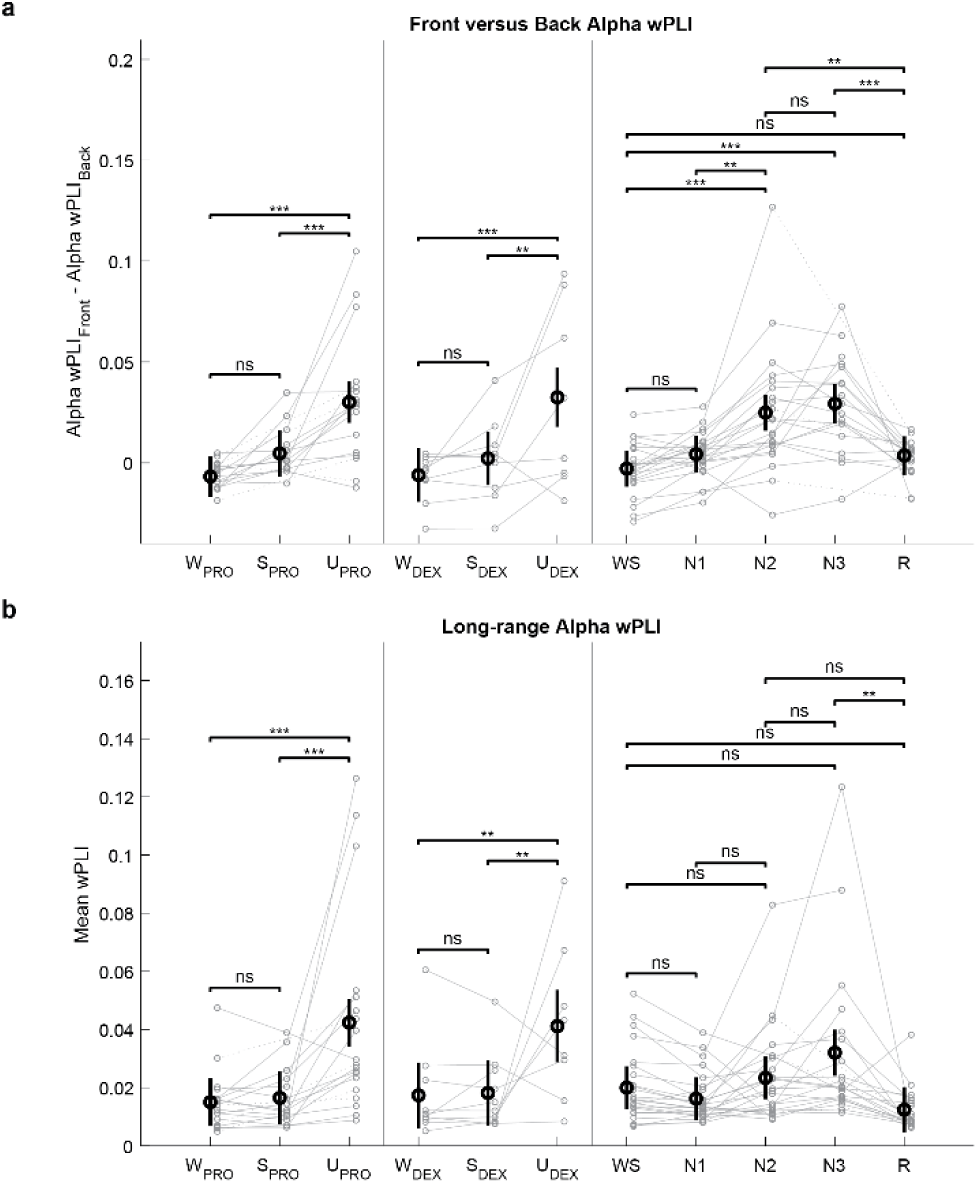
wPLI-derived outcomes. **a)** Difference in wPLI within “front” regions compared to “back” regions. Gray symbols with lines are means across recording sites for each participant; dashed lines are used when no data for an intervening behavioral state were available for that participant. Black symbols and error bars are predicted marginal means and 95% confidence intervals for the mean across participants. * *p* < 0.05, ** *p* < 0.01, *** *p* < 0.001, ns = not significant. **b)** Same as **(a)**, for average wPLI across only the longest half of connections (based on physical distance).

---

**Supplementary Table 1.** Number of segments (minutes) analyzed per participant and responsiveness state or sleep stage.

|  | Behavioral state/sleep stage |  |  |  |  |  |  |  |  |  |  | Total |
| --- | --- | --- | --- | --- | --- | --- | --- | --- | --- | --- | --- | --- |
|  | W<sub>PRO</sub> | S<sub>PRO</sub> | U<sub>PRO</sub> | W<sub>DEX</sub> | S<sub>DEX</sub> | U<sub>DEX</sub> | WS | N1 | N2 | N3 | R |  |
| Participant |  |  |  |  |  |  |  |  |  |  |  |  |
| 369R | - | - | - | - | - | - | 98 | 5 | 10 | - | - | 113 |
| 372L | 6 | 1 | 7 | - | - | - | 127 | 16 | 178 | - | 47 | 382 |
| 376R | 6 | 6 | 6 | - | - | - | 150 | 12 | 285 | 42 | 47 | 554 |
| 384B | 6 | - | 12 | - | - | - | - | - | - | - | - | 18 |
| 394R | 6 | 6 | 6 | - | - | - | - | - | - | - | - | 18 |
| 399R | 5 | 1 | 7 | - | - | - | - | - | - | - | - | 13 |
| 400L | 6 | 7 | 1 | - | - | - | - | - | - | - | - | 14 |
| 403L | 6 | 1 | 7 | - | - | - | 112 | 30 | 140 | - | 1 | 297 |
| 405L | 6 | 1 | 7 | - | - | - | - | - | - | - | - | 14 |
| 409L | 6 | 7 | 1 | - | - | - | 165 | 46 | 189 | 9 | 82 | 505 |
| 413R | 6 | 6 | 9 | - | - | - | - | - | - | - | - | 21 |
| 418R | 6 | 12 | 6 | - | - | - | 131 | 34 | 252 | 33 | 150 | 624 |
| 423L | 6 | 7 | 7 | - | - | - | 305 | 32 | 186 | 20 | 42 | 605 |
| 439B | - | - | - | 4 | 7 | 10 | 196 | 32 | 75 | 65 | - | 389 |
| 456R | - | - | - | 5 | 14 | - | - | - | - | - | - | 19 |
| 457B | - | - | - | 5 | 8 | 4 | 284 | 10 | 180 | 69 | 77 | 637 |
| 458R | - | - | - | 5 | 14 | 5 | - | - | - | - | - | 24 |
| 460L | - | - | - | 5 | 25 | - | 70 | 43 | 243 | 41 | 30 | 457 |
| 514L | 6 | - | 6 | - | - | - | 72 | 67 | 338 | 10 | 39 | 538 |
| 524R | - | - | - | - | - | - | 72 | 7 | 313 | 25 | 64 | 481 |
| 525L | - | - | - | 5 | 7 | 10 | - | - | - | - | - | 22 |
| 532R | - | - | - | - | - | - | 289 | 9 | 152 | 34 | 79 | 563 |
| 559R | - | - | - | 5 | 7 | 12 | - | - | - | - | - | 24 |
| 567R | 5 | - | 11 | - | - | - | 315 | 105 | 252 | 8 | - | 696 |
| 585L | 10 | - | 5 | - | - | - | 224 | 34 | 191 | 41 | 162 | 667 |
| 625L | - | - | - | 5 | 8 | 6 | 193 | 29 | 140 | 32 | 145 | 558 |
| 634L | 6 | 6 | 8 | - | - | - | 140 | 8 | 154 | 16 | 4 | 342 |
| 640L | 7 | 1 | 8 | - | - | - | 209 | 17 | 144 | 71 | 116 | 573 |
| 672R | 7 | 7 | 7 | - | - | - | 433 | 22 | 212 | 32 | 86 | 806 |
| 717R | - | - | - | - | - | - | 139 | 32 | 192 | 34 | 39 | 436 |
| 720R | - | - | - | 5 | 14 | 7 | 163 | 34 | 163 | 43 | 59 | 488 |
| 728R | - | - | - | 5 | 7 | 12 | 179 | 36 | 238 | 7 | 59 | 543 |
| 741L | 7 | 8 | 8 | - | - | - | 175 | 37 | 177 | - | 90 | 502 |
| 764R | - | - | - | - | - | - | 215 | 16 | 115 | 13 | 11 | 370 |
| Total | 119 | 77 | 129 | 49 | 111 | 66 | 4,456 | 713 | 4,519 | 645 | 1,429 | 12,313 |

**Supplementary Table 2.** Planned contrasts for regional delta power. Est = estimated difference. CI = confidence interval. SE = standard error. df = degrees of freedom using Satterthwaite approximation.

Frontal delta power
| Contrast | Est (dB) | 95% CI | SE | df | t ratio | p |
| --- | --- | --- | --- | --- | --- | --- |
| S<sub>PRO</sub> - W<sub>PRO</sub> | 0.7 | [-1.1, 2.5] | 0.75 | 124.5 | 0.953 | 0.61 |
| U<sub>PRO</sub> - W<sub>PRO</sub> | 4.2 | [2.6, 5.8] | 0.69 | 120.5 | 6.12 | <0.001 |
| U<sub>PRO</sub> - S<sub>PRO</sub> | 3.5 | [1.7, 5.3] | 0.75 | 124.5 | 4.62 | <0.001 |
| S<sub>DEX</sub> - W<sub>DEX</sub> | 1.1 | [-1.0, 3.2] | 0.89 | 120.5 | 1.20 | 0.45 |
| U<sub>DEX</sub> - W<sub>DEX</sub> | 5.0 | [2.7, 7.3] | 0.97 | 123.6 | 5.21 | <0.001 |
| U<sub>DEX</sub> - S<sub>DEX</sub> | 4.0 | [1.7, 6.3] | 0.97 | 123.6 | 4.10 | <0.001 |
| N1 - WS | 1.9 | [0.3, 3.5] | 0.60 | 120.5 | 3.15 | 0.014 |
| N2 - WS | 5.5 | [3.9, 7.2] | 0.60 | 120.5 | 9.17 | <0.001 |
| N3 - WS | 8.9 | [7.2, 10.6] | 0.64 | 121.5 | 13.9 | <0.001 |
| R - WS | 0.9 | [-0.8, 2.7] | 0.63 | 121.3 | 1.50 | 0.53 |
| N2 - N1 | 3.6 | [2.0, 5.3] | 0.60 | 120.5 | 6.03 | <0.001 |
| N3 - N2 | 3.4 | [1.6, 5.1] | 0.64 | 121.5 | 5.25 | <0.001 |
| R - N2 | -4.6 | [-6.3, -2.9] | 0.63 | 121.3 | -7.26 | <0.001 |
| R - N3 | -8.0 | [-9.8, -6.2] | 0.67 | 122.0 | -11.9 | <0.001 |

Parietal delta power
| Contrast | Est (dB) | 95% CI | SE | df | t ratio | p |
| --- | --- | --- | --- | --- | --- | --- |
| S<sub>PRO</sub> - W<sub>PRO</sub> | 1.5 | [0.0, 3.1] | 0.65 | 118.0 | 2.37 | 0.050 |
| U<sub>PRO</sub> - W<sub>PRO</sub> | 2.5 | [1.1, 3.9] | 0.58 | 115.5 | 4.28 | <0.001 |
| U<sub>PRO</sub> - S<sub>PRO</sub> | 1.0 | [-0.6, 2.5] | 0.65 | 118.0 | 1.48 | 0.30 |
| S<sub>DEX</sub> - W<sub>DEX</sub> | 1.0 | [-0.8, 2.7] | 0.74 | 115.5 | 1.31 | 0.39 |
| U<sub>DEX</sub> - W<sub>DEX</sub> | 5.9 | [4.0, 7.8] | 0.80 | 117.4 | 7.36 | <0.001 |
| U<sub>DEX</sub> - S<sub>DEX</sub> | 4.9 | [3.0, 6.8] | 0.80 | 117.4 | 6.15 | <0.001 |
| N1 - WS | 1.7 | [0.4, 3.1] | 0.51 | 115.5 | 3.43 | 0.0060 |
| N2 - WS | 4.9 | [3.5, 6.2] | 0.51 | 115.5 | 9.55 | <0.001 |
| N3 - WS | 8.7 | [7.3, 10.2] | 0.53 | 116.0 | 16.3 | <0.001 |
| R - WS | 1.2 | [-0.2, 2.7] | 0.53 | 116.0 | 2.31 | 0.13 |
| N2 - N1 | 3.1 | [1.7, 4.5] | 0.51 | 115.5 | 6.11 | <0.001 |
| N3 - N2 | 3.9 | [2.4, 5.3] | 0.53 | 116.0 | 7.22 | <0.001 |
| R - N2 | -3.6 | [-5.1, -2.2] | 0.53 | 116.0 | -6.77 | <0.001 |
| R - N3 | -7.5 | [-9.0, -6.0] | 0.56 | 116.2 | -13.4 | <0.001 |

Temporal delta power
| Contrast | Est (dB) | 95% CI | SE | df | t ratio | p |
| --- | --- | --- | --- | --- | --- | --- |
| S<sub>PRO</sub> - W<sub>PRO</sub> | 0.7 | [-0.9, 2.4] | 0.69 | 135.7 | 1.06 | 0.54 |
| U<sub>PRO</sub> - W<sub>PRO</sub> | 1.0 | [-0.5, 2.5] | 0.63 | 132.4 | 1.56 | 0.27 |
| U<sub>PRO</sub> - S<sub>PRO</sub> | 0.3 | [-1.4, 1.9] | 0.69 | 135.7 | 0.369 | 0.93 |
| S<sub>DEX</sub> - W<sub>DEX</sub> | 0.8 | [-1.2, 2.9] | 0.87 | 132.4 | 0.949 | 0.61 |
| U<sub>DEX</sub> - W<sub>DEX</sub> | 2.9 | [0.7, 5.2] | 0.94 | 135.3 | 3.11 | 0.0064 |
| U<sub>DEX</sub> - S<sub>DEX</sub> | 2.1 | [-0.1, 4.4] | 0.94 | 135.3 | 2.24 | 0.069 |
| N1 - WS | 1.1 | [-0.4, 2.6] | 0.56 | 132.4 | 1.95 | 0.27 |
| N2 - WS | 3.1 | [1.6, 4.7] | 0.56 | 132.4 | 5.56 | <0.001 |
| N3 - WS | 6.5 | [4.9, 8.1] | 0.60 | 133.3 | 10.9 | <0.001 |
| R - WS | 0.5 | [-1.1, 2.1] | 0.59 | 133.0 | 0.893 | 0.88 |
| N2 - N1 | 2.0 | [0.5, 3.6] | 0.56 | 132.4 | 3.61 | 0.0032 |
| N3 - N2 | 3.4 | [1.8, 5.0] | 0.60 | 133.3 | 5.65 | <0.001 |
| R - N2 | -2.6 | [-4.2, -1.0] | 0.59 | 133.0 | -4.44 | <0.001 |
| R - N3 | -6.0 | [-7.6, -4.3] | 0.62 | 133.6 | -9.66 | <0.001 |

| Contrast | Est (dB) | 95% CI | SE | df | t ratio | p |
| --- | --- | --- | --- | --- | --- | --- |
| S<sub>PRO</sub> - W<sub>PRO</sub> | 0.7 | [-0.7, 2.2] | 0.60 | 136.5 | 1.20 | 0.46 |
| U<sub>PRO</sub> - W<sub>PRO</sub> | 2.2 | [0.9, 3.5] | 0.55 | 133.2 | 3.91 | <0.001 |
| U<sub>PRO</sub> - S<sub>PRO</sub> | 1.4 | [0.0, 2.9] | 0.60 | 136.5 | 2.40 | 0.047 |
| S<sub>DEX</sub> - W<sub>DEX</sub> | 1.3 | [-0.5, 3.2] | 0.76 | 133.2 | 1.76 | 0.19 |
| U<sub>DEX</sub> - W<sub>DEX</sub> | 5.0 | [3.0, 6.9] | 0.83 | 136.2 | 5.99 | <0.001 |
| U<sub>DEX</sub> - S<sub>DEX</sub> | 3.6 | [1.6, 5.6] | 0.83 | 136.2 | 4.36 | <0.001 |
| N1 - WS | 2.0 | [0.6, 3.3] | 0.49 | 133.2 | 3.96 | <0.001 |
| N2 - WS | 4.3 | [3.0, 5.7] | 0.49 | 133.2 | 8.80 | <0.001 |
| N3 - WS | 7.2 | [5.8, 8.7] | 0.52 | 134.1 | 13.8 | <0.001 |
| R - WS | 0.9 | [-0.5, 2.2] | 0.52 | 133.9 | 1.65 | 0.43 |
| N2 - N1 | 2.4 | [1.1, 3.7] | 0.49 | 133.2 | 4.83 | <0.001 |
| N3 - N2 | 2.9 | [1.5, 4.3] | 0.52 | 134.1 | 5.54 | <0.001 |
| R - N2 | -3.5 | [-4.9, -2.1] | 0.52 | 133.9 | -6.78 | <0.001 |
| R - N3 | -6.4 | [-7.9, -4.9] | 0.54 | 134.4 | -11.8 | <0.001 |

**Supplementary Table 3.**
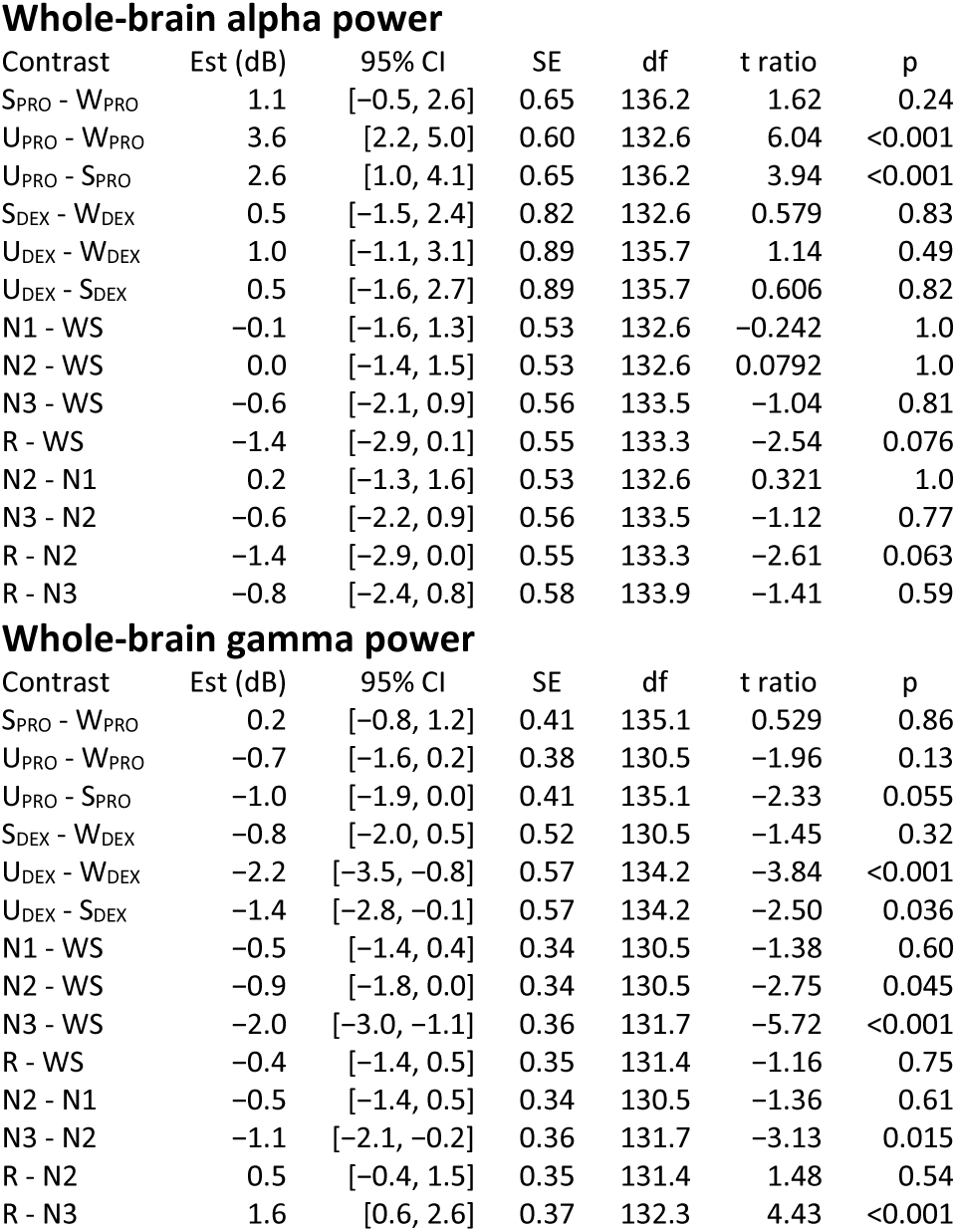
Planned contrasts for whole-brain alpha and gamma power. Est = estimated difference. CI = confidence interval. SE = standard error. df = degrees of freedom using Satterthwaite approximation.

**Supplementary Table 4.** Planned contrasts for frontal alpha power. Est = estimated difference. CI = confidence interval. SE = standard error. df = degrees of freedom using Satterthwaite approximation.

| Contrast | Est (dB) | 95% CI | SE | df | t ratio | p |
| --- | --- | --- | --- | --- | --- | --- |
| S<sub>PRO</sub> - W<sub>PRO</sub> | 1.9 | [-0.1, 3.9] | 0.85 | 124.6 | 2.26 | 0.065 |
| U<sub>PRO</sub> - W<sub>PRO</sub> | 8.0 | [6.1, 9.8] | 0.77 | 119.6 | 10.3 | <0.001 |
| U<sub>PRO</sub> - S<sub>PRO</sub> | 6.0 | [4.0, 8.1] | 0.85 | 124.6 | 7.15 | <0.001 |
| S<sub>DEX</sub> - W<sub>DEX</sub> | 0.8 | [-1.6, 3.2] | 1.00 | 119.6 | 0.764 | 0.73 |
| U<sub>DEX</sub> - W<sub>DEX</sub> | 2.8 | [0.3, 5.4] | 1.09 | 123.4 | 2.61 | 0.027 |
| U<sub>DEX</sub> - S<sub>DEX</sub> | 2.1 | [-0.5, 4.6] | 1.09 | 123.4 | 1.91 | 0.14 |
| N1 - WS | 0.3 | [-1.5, 2.1] | 0.68 | 119.6 | 0.441 | 0.99 |
| N2 - WS | 1.4 | [-0.4, 3.3] | 0.68 | 119.6 | 2.12 | 0.19 |
| N3 - WS | 1.3 | [-0.6, 3.3] | 0.72 | 121.0 | 1.84 | 0.32 |
| R - WS | -0.8 | [-2.7, 1.2] | 0.71 | 120.6 | -1.06 | 0.80 |
| N2 - N1 | 1.1 | [-0.7, 3.0] | 0.68 | 119.6 | 1.68 | 0.41 |
| N3 - N2 | -0.1 | [-2.1, 1.8] | 0.72 | 121.0 | -0.150 | 1.0 |
| R - N2 | -2.2 | [-4.1, -0.3] | 0.71 | 120.6 | -3.09 | 0.018 |
| R - N3 | -2.1 | [-4.1, -0.1] | 0.75 | 121.6 | -2.78 | 0.041 |

**Supplementary Table 5.**
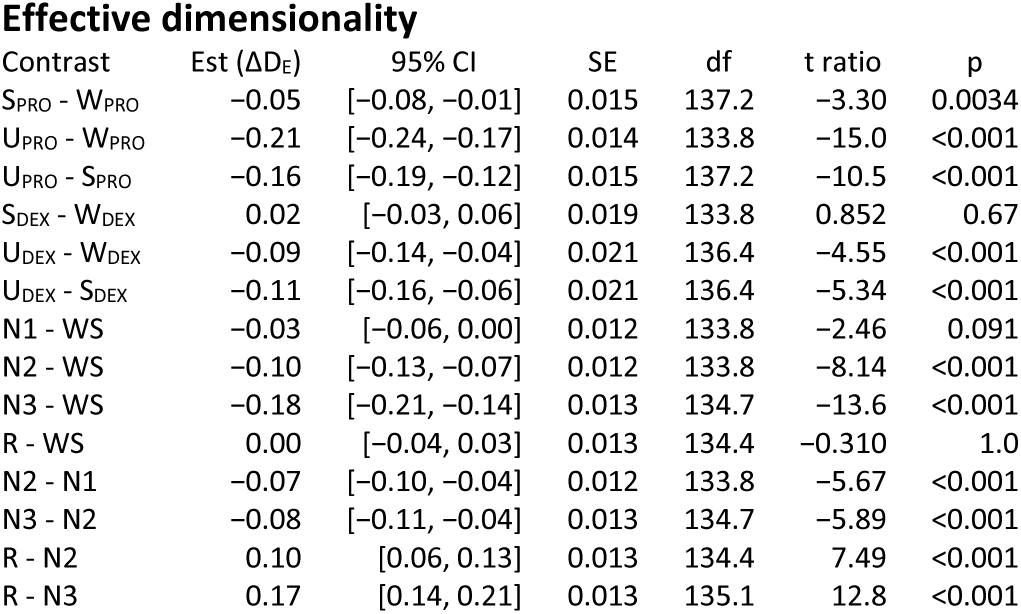
Planned contrasts for effective dimensionality. Est = estimated difference. CI = confidence interval. SE = standard error. df = degrees of freedom using Satterthwaite approximation.

**Supplementary Table 6.** Planned contrasts for alpha wPLI outcomes. Est = estimated difference. CI = confidence interval. SE = standard error. df = degrees of freedom using Satterthwaite approximation.

Alpha wPLI Front minus Back
| Contrast | Est ($\Delta wPLI$) | 95% CI | SE | df | t ratio | p |
| --- | --- | --- | --- | --- | --- | --- |
| S<sub>PRO</sub> - W<sub>PRO</sub> | 0.012 | [-0.004, 0.027] | 0.007 | 127.1 | 1.72 | 0.20 |
| U<sub>PRO</sub> - W<sub>PRO</sub> | 0.037 | [0.022, 0.051] | 0.006 | 120.1 | 5.98 | <0.001 |
| U<sub>PRO</sub> - S<sub>PRO</sub> | 0.025 | [0.009, 0.041] | 0.007 | 127.1 | 3.77 | <0.001 |
| S<sub>DEX</sub> - W<sub>DEX</sub> | 0.008 | [-0.011, 0.027] | 0.008 | 120.1 | 1.03 | 0.56 |
| U<sub>DEX</sub> - W<sub>DEX</sub> | 0.039 | [0.018, 0.059] | 0.009 | 125.9 | 4.46 | <0.001 |
| U<sub>DEX</sub> - S<sub>DEX</sub> | 0.030 | [0.010, 0.051] | 0.009 | 125.9 | 3.51 | 0.0019 |
| N1 - WS | 0.007 | [-0.008, 0.022] | 0.005 | 120.1 | 1.32 | 0.65 |
| N2 - WS | 0.028 | [0.013, 0.042] | 0.005 | 120.1 | 5.12 | <0.001 |
| N3 - WS | 0.032 | [0.017, 0.048] | 0.006 | 122.9 | 5.59 | <0.001 |
| R - WS | 0.006 | [-0.009, 0.022] | 0.006 | 122.2 | 1.15 | 0.75 |
| N2 - N1 | 0.021 | [0.006, 0.035] | 0.005 | 120.1 | 3.81 | 0.0016 |
| N3 - N2 | 0.004 | [-0.011, 0.020] | 0.006 | 122.9 | 0.764 | 0.93 |
| R - N2 | -0.021 | [-0.037, -0.006] | 0.006 | 122.2 | -3.76 | 0.0020 |
| R - N3 | -0.026 | [-0.042, -0.010] | 0.006 | 123.7 | -4.30 | <0.001 |

| Contrast | Est ($\Delta wPLI$) | 95% CI | SE | df | t ratio | p |
| --- | --- | --- | --- | --- | --- | --- |
| S<sub>PRO</sub> - W<sub>PRO</sub> | 0.001 | [-0.012, 0.015] | 0.006 | 140.0 | 0.266 | 0.96 |
| U<sub>PRO</sub> - W<sub>PRO</sub> | 0.027 | [0.015, 0.040] | 0.005 | 132.1 | 5.22 | <0.001 |
| U<sub>PRO</sub> - S<sub>PRO</sub> | 0.026 | [0.013, 0.039] | 0.006 | 140.0 | 4.59 | <0.001 |
| S<sub>DEX</sub> - W<sub>DEX</sub> | 0.001 | [-0.016, 0.018] | 0.007 | 132.1 | 0.109 | 0.99 |
| U<sub>DEX</sub> - W<sub>DEX</sub> | 0.024 | [0.005, 0.042] | 0.008 | 139.4 | 3.07 | 0.0072 |
| U<sub>DEX</sub> - S<sub>DEX</sub> | 0.023 | [0.005, 0.041] | 0.008 | 139.4 | 2.97 | 0.0097 |
| N1 - WS | -0.004 | [-0.016, 0.009] | 0.005 | 132.1 | -0.805 | 0.92 |
| N2 - WS | 0.003 | [-0.009, 0.016] | 0.005 | 132.1 | 0.722 | 0.94 |
| N3 - WS | 0.012 | [-0.001, 0.025] | 0.005 | 135.7 | 2.45 | 0.094 |
| R - WS | -0.008 | [-0.021, 0.006] | 0.005 | 134.8 | -1.56 | 0.48 |
| N2 - N1 | 0.007 | [-0.005, 0.020] | 0.005 | 132.1 | 1.53 | 0.51 |
| N3 - N2 | 0.009 | [-0.005, 0.022] | 0.005 | 135.7 | 1.76 | 0.36 |
| R - N2 | -0.011 | [-0.024, 0.002] | 0.005 | 134.8 | -2.26 | 0.14 |
| R - N3 | -0.020 | [-0.033, -0.006] | 0.005 | 136.9 | -3.86 | 0.0013 |

